# Seasonal changes of metabolites in phloem sap from *Broussonetia papyrifera*

**DOI:** 10.1101/317271

**Authors:** Jiang-tao Shi, Hai-chong Liu, Jia-yan Luo, Li-ping Cai

## Abstract

Gas chromatography-Mass spectrometry (GC-MS) were employed to analyze the whole metabolites in phloem sap of *Broussonetia papyrifera* and the seasonal changes of content of these metabolites were also investigated. Thirty-eight metabolites were detected in *BP* phloem exudates. The highest content (44.59mg g^-1^) of total metabolites was presented in March. High contents of organic acids and sugars were detected in *BP* phloem exudates from all growing months. Smaller amounts of fatty acids and alcohols were also detected in BP phloem exudates. Interestingly, some metabolites, such as PI3 kinase inhibitor, Chlorogenic acid, Chelerythrine and palmitic acid, which have properties of bioactivity to anticancer and anti-inflammation, were also detected. Quininic acid was the most abundant organic acid, representing up to 86.3% (average value) of all organic acids. D-fructose, D-glucose, and sucrose were the major soluble sugars in phloem saps and the maximum of sugars content was 19.76mg g^-1^ (average value) in November. Seasonal changes of contents of metabolites were different among individuals. The metabolites analysis double confirmed that the *BP* phloem sap can be serviced as an important resource for synthesis of pharmaceutical and human health products.

## Introduction

Well-known as its fast growing, good adaptability, strong germinating ability and outstanding regeneration capability, *Broussonetia papyrifera* L. Vent. (BP) is widely planted in East Asia, the United States, and the Pacific Islands[1,2]. Similar to other tree species from Moraceae, *BP* has more abundant sap from its phloem tissue. Previous studies reported that some bioactive ingredients were extracted from the *BP* phloem sap and they were proved to be beneficial to antioxidant[3], anti-inflammatory and anticancer[4–6] and shown inhibitory activity against aromatase[7], tyrosinase[8], α-Glucosidase[9], and anticholinesterase[10]. Consequently, the *BP* phloem sap is a promising natural resource for synthesis of new medicines and health products. Otherwise, *BP* phole is a good choice for pulping and manufacturing of high quality paper[11,12]. To understand the chemical composition in phloem sap is beneficial to improvement of paper industrial.

Phloem sap is produced from the secondary metabolites of tree and relative to tree physiology. Fresh phloem sap commonly consists of polychemical compositions, which is produced from metabolic active of photosynthate. These chemical ingredients play key roles on tree physiology, such as resistance of pest infestation [13,14], maintenance of osmotic pressure between conductivity cells[15,16], and adjustment of the balance of inorganic ions[16]. Metabolic profiling is an effective technology for investigating the chemical composition in plant tissue[17–21]. However, limited studies focusing on the metabolites and their seasonal changes of *BP* phloem sap were found. As well-known, tree physiology is regulated by endogenous and exogenous factors. As the most important exogenous regulatory factors, the ttemperature, illumination and precipitation changes in different growing months. In addition, the phloem sap is a secretion from laticifer cells in tree phloem tissue, which is divided yearly from cambium[22]. Many studies have proved that the anatomy structure in phloem formation shows a seasonal dynamic change during growing years[23–29]. Therefore, it was hypothesized that there were seasonal changes on the chemical compositions and their concentration of the *BP* phloem sap. Based on the mentioned previously, the aim of this work was to investigate the chemical compositions in *BP* phloem sap by GC-MS and intended to reveal the seasonal changes of these compounds in Nanjing city of China during the 2016 growing season.

## Experimental

### Phloem sap collection

Five healthy *Broussonetia papyrifera* trees were selected from campus of Nanjing Forestry University (NJFU) (32°04’57.4’’ N, 118°49’00.3’’ E), Nanjing, China. The average dimeter at breast height was 17.6 cm and the average tree age was 11.5 years. Phloem saps were collected between 10 a.m. to 12 a.m. at Mar. 28, Apr. 22, May 20, Jun.15, Jul.22, Sep.20, Oct.21, and Nov.15. Sampling of phloem saps was simple and feasible because phloem saps are mainly stored in axial laticifer cells. The cell types in *BP* phloem is shown in Fig 1. *BP* phloem generally contained sieve tubes, laticifers, phloem rays, phloem fibers, and crystals. Even though in the region of nonconducting phloem featured with collapsed sieve tubes, latidifer cells still scatter intact[22,30,31]. Briefly, after peeling off the outset bark (approximately 2 cm^2^), the phloem was gashed by disinfected scalpel at a 45° angle to tree growth direction. The saps were natural flowed down into prepared tubes and frozen by liquid *N*_2_ immediately and stored in −20°C for future use. In order to avoid the interaction on saps from each month, sampling position on tree truck was spiral and interval 10 cm between each other at all directions.

**Fig.1.**
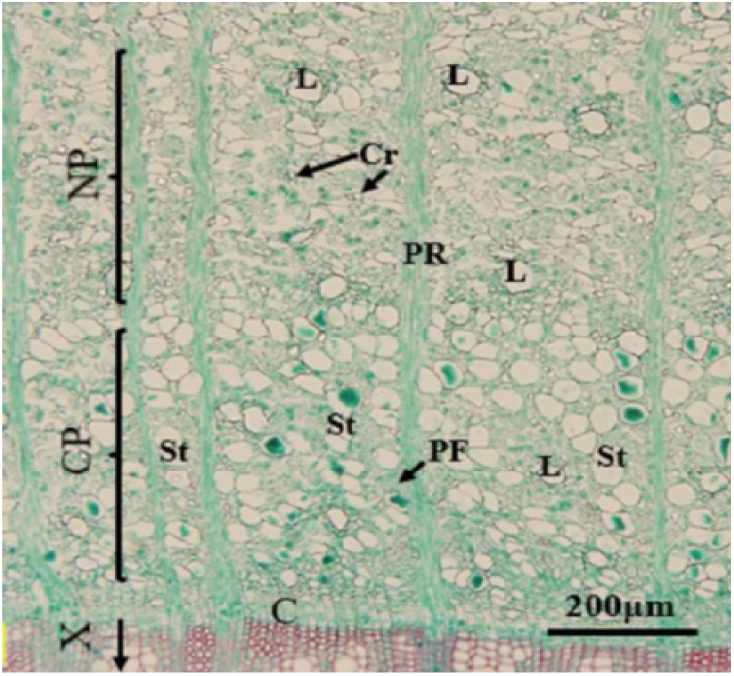
Cross section and cells in *BP* phloem collected in October, 2016. C: cambium; X: xylem; L: laticifers; CP: conducting phloem; NP: nonconducting phloem; Cr: crystals; PR: phloem rays; PF: phloem fibers; St: sieve tubes. Bar is 200μm.

### Pre-treatment of the sap

Phloem saps were extracted by the mixture solution of methanol, chloroform and deionized water according to the former reports[19,21]. The freeze saps were finely grinded under liquid N2 condition and 50 ± 3mg sample was mixed with 1mL methanol (-20°C pre-chilled) in 1.5mL centrifuge tube. Then 50μl ribitol (CAS No.488-81-3, 2mg mL^-1^ in ddH_2_O), which was selected as internal standard for compounds quantitative analysis, was added and the mixture was incubated for 20min at 70°C and 950rpm min^-1^. Following by 15min centrifuge at 10000rpm min^-1^, 500μl supernatant was transferred to a new tube and mixed with equal volume of chloroform (-20°C pre-chilled) and deionized water (4°C pre-chilled), and then incubated for 20min at 37°C at and 950rpm min^-1^. After 15min centrifuging at 4000rpm min^-1^, about 500μl supernatant was transferred to a new tube and stored at −20°C for further test. Ribitol and extraction reagents were purchased from Sigma-Aldrich (USA) and Nanjing chemical reagent Co., Ltd, China, respectively.

### Derivatization and GC-MS

The frozen pure phloem sap was dried under vacuum at −60°C and derivatized with *N*-methyl-*N*-trimethlsilyl-trifluoroacetamide (MSTFA) (CAS No.24589-78-4). Briefly, the dried samples were mixed with 50μl Methoxyamine hydrochloride (CAS No.593-56-6, dissolved in pyridine, CAS No.110-86-1, 20mg ml^-1^) and incubated 2 hours at 37°C, 950rpm min^-1^. Then 100μl MSTFA (containing 20μl ml^-1^ alkanes) was added and kept 30 min at 37°C, 950rpm min^-1^ and overnight at room temperature. All the derivatization reagents were purchased from Sigma-Aldrich (USA). 0.4μL derivatized samples were injected into GC-MS (TRACE DSQ, USA) fitted with a DB-5MS column (30m ×0.32 mm×0.25μm). The flow rate for the He (99.999%) carrier gas was 1.0mL min^-1^. The detail parameters for GC test were as follows: the initial temperature was held at 50°C for 1min, and then increased to 300°C at a rate of 10°C min^-1^, held for 5min. Both the injector and the detector temperatures were 250°C. EI ion and ionization voltage was pre-set at 70 eV and scan range was 20-500 amu.

### MS peak identification and data analysis

The Gas chromatogram was observed using the MS Workstation version 6.9.3 and the online result was searched with the NIST (National Institute of Standard and Technology, USA) mass spectrum data, and referred to related literature for metabolites identification and classification[17,20,21]. The relative content of metabolites was calculated using the equation of C_x_=[(A_x_/A_i_)×0.05×2mg mL^-1^]/m_0_ (mg g^-1^), where C_x_ is the relative content of identified metabolite; A_x_ is the peak area of identified metabolite; A_i_ is the peak area of internal standard; m_0_ is the dry weight of phloem tissue[18,21]. Excel 2016 was used to draw the charts.

## Results and discussion

### Chemical composition in phloem sap of BP

Metabolites of the exudates from the *BP* phloem were analyzed by GC-MS after being derivatized by MSTFA. Thirty-eight metabolites were identified and their concentration was calculated by the mass of each metabolite with drying weight of phloem tissue. The mean values of the total concentration of metabolites and the percentage composition of *BP* phloem sap are shown in Fig. 2. The Highest (44.59mg g^-1^) concentration of total metabolites occurred in March and the total concentration decreased to the lowest value (6.78mg g^-1^) in May, then increased to 23.01 mg g^-1^ in June. In July, the total concentration of metabolites was 9.59mg g^-1^, which was the secondary lowest during the whole growing season. The total concentration increased to 20.96 mg g^-1^ in September and decreased slightly in October. The Phloem sap collected in November had the secondary highest concentration of total metabolites.

**Fig.2.**
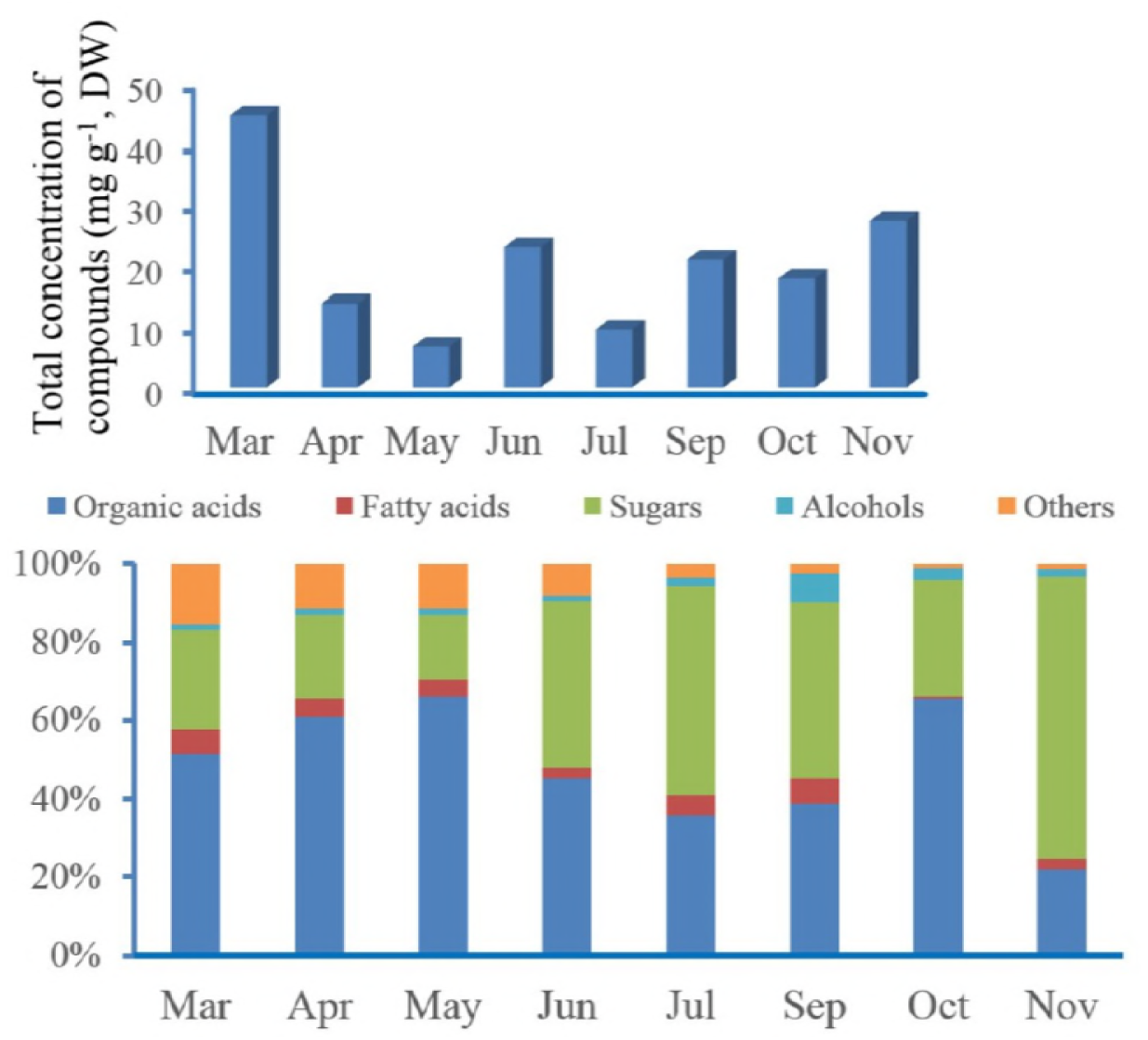
Seasonal variation of total concentration of metabolites in *BP* phloem sap and the percentage of composition of the major groups. Mean value was used for drawing.

As shown in Fig.2, all identificated metabolites can be classified into organic acids, sugars, fatty acids and alcohols. In all growing months, organic acids and sugars represented on the average of 86.6% of the overall metabolite concentration of the *BP* phloem sap. The Previous work stated that a wide range of sugars and organic acids can be detect by the MSTFA derivatized method [32]. Fatty acids and alcohols were abundant compositions of the *BP* phloem sap exudates, both of them representing 1%-7% of the total metabolites during all growing months. Other metabolites such as PI3-Kinase Inhibitor, silanamine, were also identified, although they represented less than 1% of the total. The changes of percentages of different metabolites depended on the growing month. The Highest percentage of organic acid presented in May (66% of the total metabolites) and lowest percentage existed in November (22% of the total metabolites). On the contrary, the lowest percentage of sugars presented in May (17% of the total metabolites) and the highest percentage existed in November (72% of the total metabolites). Both the highest percentage of fatty acids and alcohols occurred in September (7% of the total metabolites).

### Seasonal changes of all metabolites

To estimate the seasonal variation of all metabolites in *BP* phloem sap, five trees were selected for sampling monthly. Qualitative and quantitative analysis of all identified metabolites performed on GC-MS. The chemical composition and their content changes during whole growing season 2016 are shown in Table 1 and Table 2. As mentioned previously, metabolites were discussed as two groups, i.e., organic acids and sugars, as well as others.

**Table 1.** Organic acids detected in *BP* phloem sap and the season changes of their contents (n=5). Unit: mg g^-1^

| Compounds | Mar | Apr | May | Jun | Jul | Sep | Oct | Nov |
| --- | --- | --- | --- | --- | --- | --- | --- | --- |
| *Lactic acid* | 0.559±0.062 | 0.128±0.016 | 0.068±0.008 | 0.052±0.005 | 0.031±0.007 | 0.230±0.039 | 0.447±0.053 | 0.078±0.012 |
| *Oxalic acid* | 0.157±0.032 | --- | 0.049±0.006 | --- | 0.025±0.002 | 0.120±0.039 | 0.079±0.018 | 0.049±0.013 |
| *Glycolic acid* | 0.144±0.029 | 0.054±0.011 | 0.041±0.004 | 0.057±0.011 | --- | 0.016±0.001 | 0.034±0.010 | --- |
| *Propanedioic acid* | 0.079±0.018 | 0.078±0.025 | 0.050±0.003 | 0.191±0.042 | --- | 0.024±0.003 | --- | --- |
| *3-Hydroxyisovaleric acid* | 0.392±0.063 | 0.159±0.047 | 0.095±0.009 | 0.130±0.033 | --- | --- | --- | --- |
| *Butanedioic acid* | 0.192±0.025 | --- | 0.126±0.023 | 0.052±0.007 | --- | --- | --- | --- |
| *Benzoic acid* | --- | --- | 0.018±0.000 | --- | --- | 0.030±0.002 | 0.037±0.009 | --- |
| *2-Butenedioic acid* | --- | --- | 0.215±0.048 | 0.085±0.016 | --- | --- | --- | --- |
| *Malic acid* | 0.486±0.071 | 0.070±0.002 | 0.072±0.012 | 0.105±0.027 | 0.019±0.000 | --- | 0.041±0.016 | --- |
| *Citric acid* | 0.235±0.039 | 0.353±0.039 | 0.121±0.036 | 0.575±0.055 | --- | 0.110±0.035 | --- | --- |
| *Ribonic acid* | --- | 0.190±0.022 | 0.048±0.009 | 0.239±0.038 | 0.050±0.004 | --- | --- | --- |
| *Shikimic acid* | 0.156±0.018 | --- | 0.217±0.059 | --- | --- | --- | --- | --- |
| *Quinic acid* | 19.634±2.336 | 6.915±0.319 | 3.106±0.116 | 7.575±0.407 | 3.148±0.085 | 7.576±0.526 | 10.842±1.338 | 5.662±0.527 |
| *L-Threonic acid* | 0.094±0.015 | 0.069±0.025 | 0.038±0.004 | 0.219±0.036 | --- | --- | --- | --- |
| *5-O-Feruloylquinic acid* | 0.317±0.068 | --- | 0.117±0.020 | 0.447±0.052 | --- | --- | --- | 0.035±0.005 |
| *Galactaric acid* | 0.534±0.092 | 0.398±0.051 | 0.095±0.018 | 0.737±0.049 | 0.153±0.028 | --- | 0.206±0.045 | 0.152±0.033 |
| **Total (Mean value)** | 22.980 | 8.416 | 4.478 | 10.467 | 3.426 | 8.107 | 11.687 | 5.977 |

All values were calculated by dividing the concentration of each metabolites by the dry weight of phloem tissue. Values are given as mean ±SD (n=5). Not detected is showed as “---”.

**Table 2.**
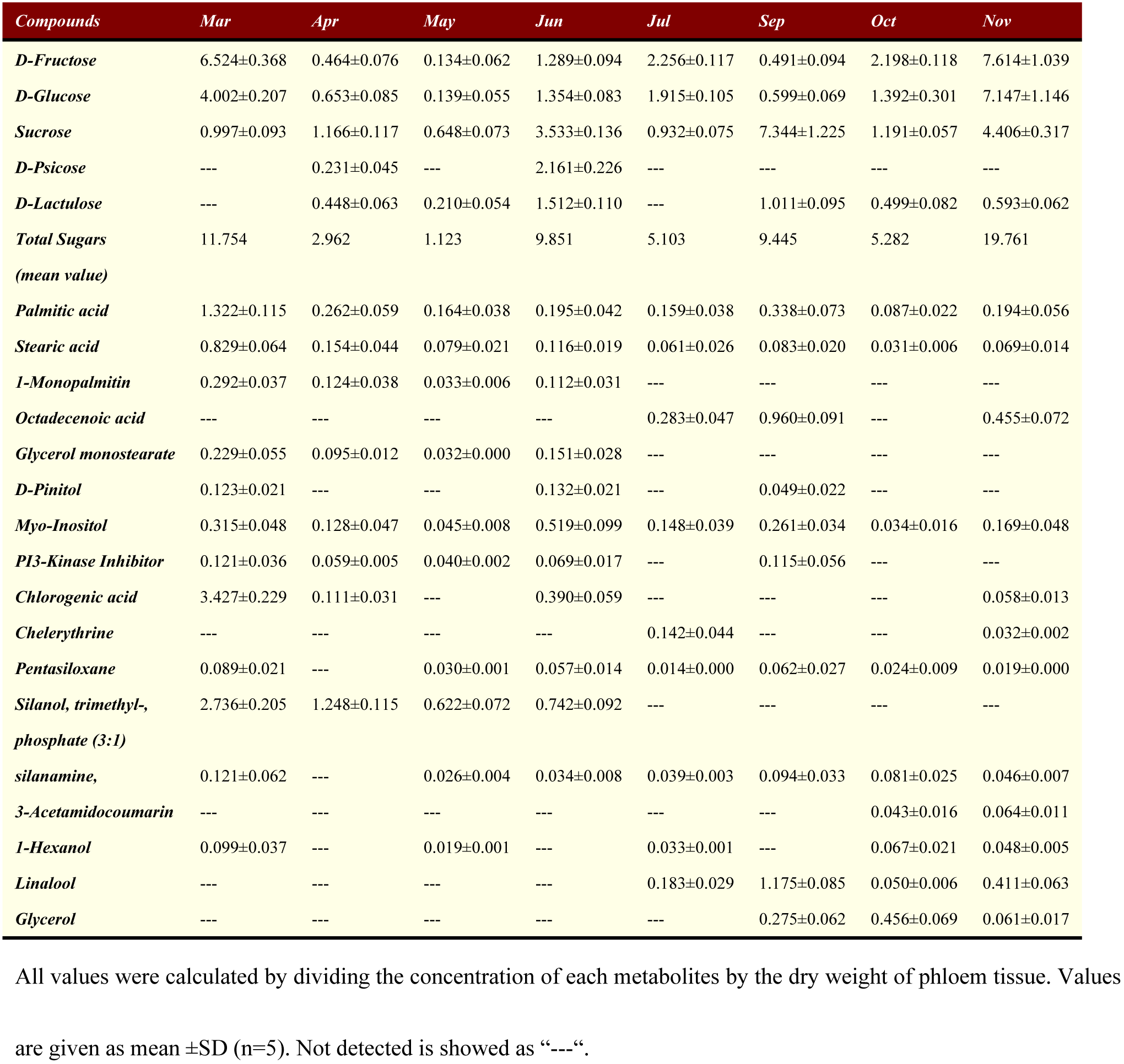
Sugars and other metabolites detected in *BP* phloem sap and their season changes (n=5). Unit: mg g^-1^

#### Organic acids

As shown in Table 1, sixteen organic acids were detected in the *BP* phloem sap and their compositions and contents changed depending on growing month. In all months, the quininic acid was the most abundant organic acid, representing up to 86.3% (average value) of all organic acids. It was showed the highest content (19.643mg g^-1^) of quininic acid at March (Table 1). But the highest percentage of quininic acid presented in November (Fig. 3). The seasonal changes of content of quininic acid were similar to that in total concentration of all metabolites. Quininic acid, which was used as an astringent and starting material for the synthesis of new pharmaceuticals, was also purified from *Eucalyptus globulus* [33], cinchona bark, and other plant products. Galactaric acid, a precursor for synthesis of adipic acid[34], was the secondary abundant organic acid in the *BP* phloem sap and its average percentage was 3.6% of all organic acid contents. Both the highest content (0.737mg g^-1^) and percentage (7.1%) of galactaric acid presented in June. But it was not detected in September. Citric acid was another abundant organic acid in the *BP* phloem sap and its highest content and percentage were 0.575 mg g^-1^ and 5.6%, respectively, in June. However, citric acid was not detected in July, October and November (Table 1). Citric acid, an important industrial organic acid, is widely used as an acidifier, flavoring and chelating agent, and is manufactured more than million tons every year[35]. On the other hand, the seasonal difference of the citric acid content is because it is an intermediate in the tricarboxylic acid cycle (TCA)[36], which is most related to the tree physiology in different growing months. Lactic acid was detected in all growing months and its highest content was 0.559mg g^-1^ in March, but the percentage of lactic acid in all organic acids was the highest in October. Lactic acid is mainly employed in food additives, pharmaceutical and cosmetics. Some other organic acids, such as oxalic acid, glycolic acid, propanedioic acid and so forth, were detected, but all of their percentages were less than 2% of total organic acids and some of them only existed in specific growing months (Table 1).

**Fig.3.**
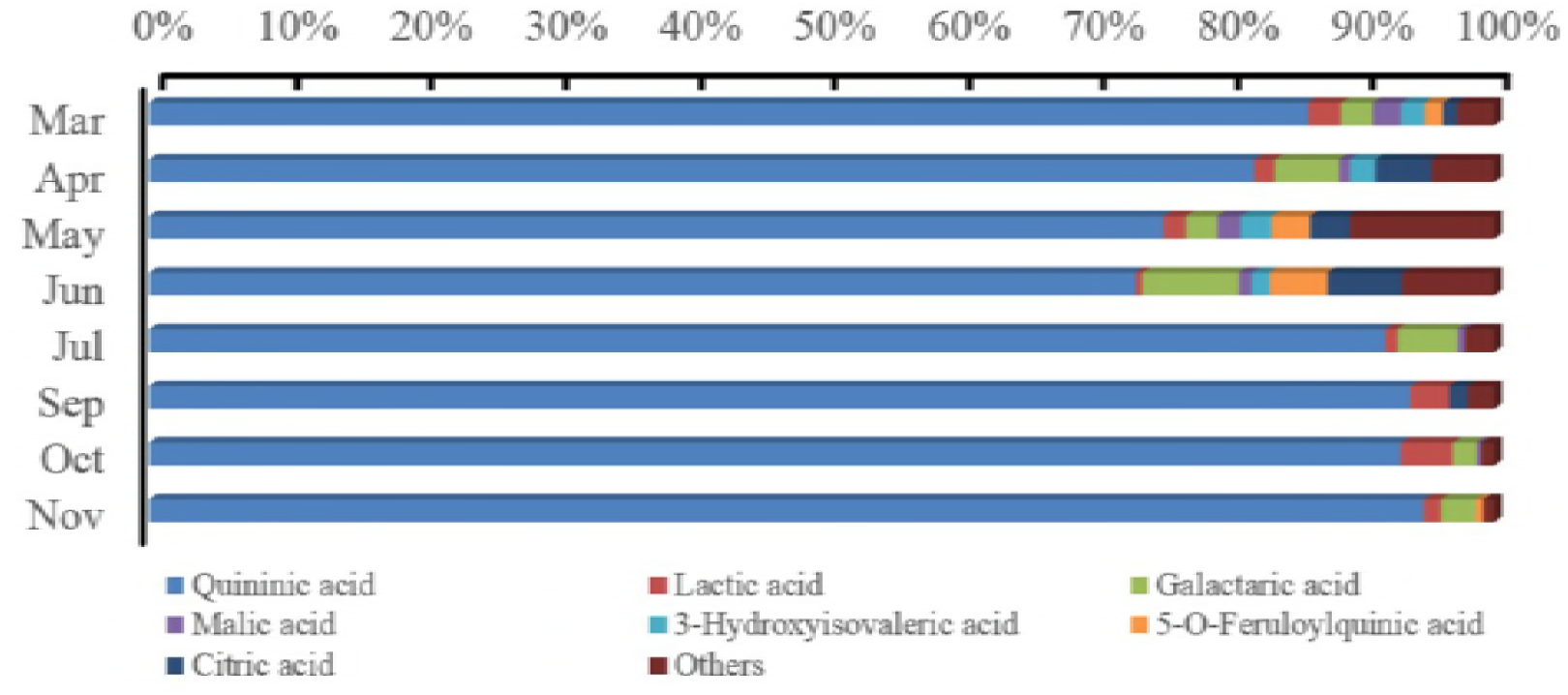
Percentage composition of organic acids in the *BP* phloem saps and their season changes.

#### Sugars and other metabolites

Natural sugars are the fundamental metabolites from photosynthesis in tree growth and provide energy and precursor for tree physiology and biosynthesis of other organic compounds. Sucrose, D-fructose, D-glucose, D-psicose, and D-lactulose were detected in the *BP* phloem sap. The highest content of total sugars was 19.761mg g^-1^ (average value) in November and the lowest was only 1.123mg g^-1^ (average value) in May (Table 2). The content and percentage of individual sugar was depending on different growing months. D-fructose was the most abundant sugar, representing almost over 40% of the soluble sugars in March, July, October, and November (Fig. 4). On the other hand, sucrose was the most abundant sugar in other four growing months. D-fructose content showed the maximum (7.614mg g^-1^) and minimum (0.134mg g^-1^) (average value) in November and May, respectively. Similarly, the highest content of D-glucose presented in November and the lowest value in May. Phloem sap collected from September contained the highest content of sucrose (7.344mg g^-1^, average value) and nearly occurring to 80% of all sugars in this month (Table 2 and Fig.4). According to previous studies, sucrose can be converted into glucose and fructose by sucrose synthase[37]. In addition, more abundant of sugars in phloem is the reason for phloem sap-sucking insects[13,14].

**Fig.4.**
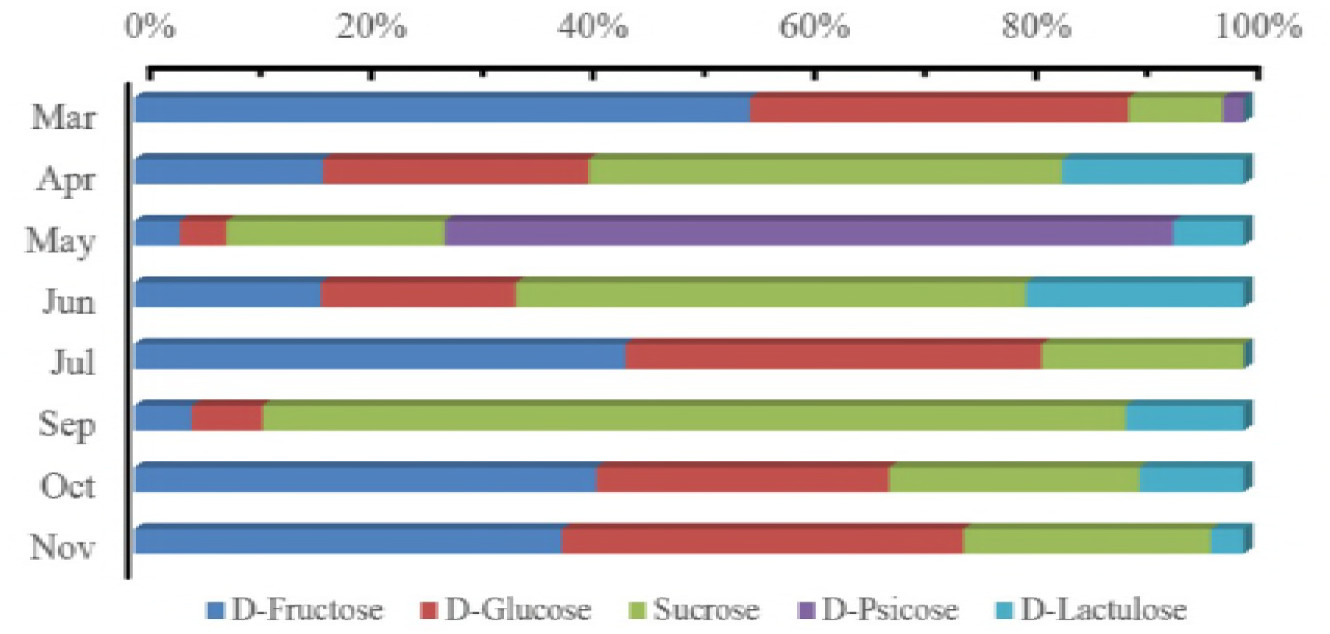
Percentage composition of sugars in the *BP* phloem saps and their season changes.

Two fatty acids, palmitic acid and stearic acid, were detected in the *BP* phloem sap in all growing months (Table 1). Moreover, it was shown the same seasonal change patterns. The highest content of both of them presented in March. Pascual et al. (2017) reported that the palmitic acid strongly boosted the metastatic potential of CD36+ in mouse models of human oral cancer cells[38]. Stearic acid is chemical industrial resource and wildly used in the production of cosmetics and lubricants. In addition, glycerol monostearate, 1-monopalmitin, and octadecenoic acid were detected in the *BP* phloem sap from partly growing months. Glycerol monostearate, a food additive and control release agent in pharmaceuticals, was also detected in the *BP* phloem sap, especially in March (0.229mg g^-1^) and June (0.151mg g^-1^) (average value).

Alcohols also are an important substrate and intermediate products in cycle of metabolism. Myo-inositol, an important intermediate in plants, was proved to have positive physiological effects on humans[39,40]. It was shown the highest content (0.519mg g^-1^, average value) of myo-inositol in June and there was distinct various in different months. Low content of D-pinitol was detected in March, June, and September. Many studies reported that the D-pinitol was a bioactive ingredient in biological and medical activities[41,42].

Interestingly, PI3 kinase inhibitor, a type of medical drug for anticancer and anti-inflammation[43–46], was detected in the *BP* phloem sap. Compared to other months, the higher contents of PI3K inhibitor were 0.121mg g^-1^ and 0.115mg g^-1^ (average values) in March and September, respectively. Chlorogenic acid, another potential medical drug in anti-inflammatory and reduction in blood pressure[47,48], was also detected in the *BP* phloem sap and its maximum was 3.427mg g^-1^ in March (Table 2). Linalool content showed the highest value of 0.411mg g^-1^ in November, which was proved to be an antifungal activity substance[49]. Chelerythrine, a benzophenanthridine alkaloids with potent anti-inflammatory effects in vivo[50], was 0.142mg g^-1^ in July.

## Conclusions

Chemical composition of the exudates from *BP* phloem were analyzed by GC-MS after being derivatized using MSTFA. Thirty-eight metabolites were detected, and the total concentration of these metabolites were different depending on growing month. The highest total metabolites concentration was 44.59mg g^-1^ in March and the lowest one was 6.78mg g^-1^ in May. Organic acids and sugars were the most abundant metabolites in the *BP* phloem sap from whole growing months. Small amounts of fatty acids and alcohols were also detected in *BP* phloem exudates. Seasonal changes of the content of metabolites were different among individuals. The highest content of organic acids was 22.98mg g^-1^ (average value) in March and quininic acid was the most abundant organic acid, representing up to 86.3% (average value) of all organic acids. In November, it was shown the maximum of sugars content of 19.76 mg g^-1^ (average value) and D-fructose, D-glucose, and sucrose were the major soluble sugars. Interestingly, some metabolites, such as PI3 kinase inhibitor, Chlorogenic acid, Chelerythrine, and palmitic acid, which have bioactivity to anticancer and anti-inflammation, were also detected and showed different seasonal variations. The metabolite analysis double confirmed that the *BP* phloem sap can be serviced as an important resource for synthesis of pharmaceutical and human health products.

## Acknowledgements

All the authors would like to thank the supporting of the National Natural Science Foundation of China (No. 31600454), Jiangsu Overseas Visiting Scholar Program for University Prominent Young & Middle-aged Teachers and Presidents and PAPD.

## Author’s contribution

Conceived and designed the experiments: JT Shi, JY Luo.

Performed the experiments: HC Liu, JT Shi.

Analyzed the data: JT Shi, HC Liu, JY Luo, LP Cai.

Wrote the manuscript and approved the final version: JT Shi, HC Liu, JY Luo, LP Cai.

## References

[1] Fahrney K, Boonnaphol O, Keoboulapha B (1997) Indigenous management of paper mulberry (*Broussonetia papyrifera*) in swidden rice fields and fallows in northern Laos. In: Paper presented at the regional workshop on indigenous strategies for intensification of shifting cultivation in Southeast Asia. Bogor, Indonesia, pp. 23–27

[2] Saito K, Linquist B, Keobualapha B et al (2009) *Broussonetia papyrifera* (paper mulberry): its growth, yield and potential as a fallow crop in slash-and-burn upland rice system of northern Laos. Agroforest Syst 3: 525–532

[3] Xu ML, Wang L, Hu JH et al (2010) Antioxidant activities and related polyphenolic constituents of the methanol extract fractions from *Broussonetia papyrifera* stem bark and wood. Food Sci Biotechnol 19(3): 677–682

[4] Wang L, Son HJ, Xu ML et al (2010) Anti-inflammatory and anticancer properties of dichloromethane and butanol fractions from the stem bark of *Broussonetia papyrifera*. J Korean Soc Appl Biol Chem 53(3): 297–303

[5] Guo FJ, Feng L, Huang C et al (2013) Prenylfavone derivatives from *Broussonetia papyrifera*, inhibit the growth of breast cancer cells in vitro and in vivo. Phytochemistry Lett 6: 331–336

[6] Park S, Fudhaili A, Oh SS et al (2016) Cytotoxic effects of kazinol A derived from *Broussonetia papyrifera* on human bladder cancer cells, T24 and T24R2. Phytomed 23: 1462–1468

[7] Lee DJ, Bhat KPL, Fong HHS et al (2001) Aromatase inhibitors from *Broussonetia papyrifera*. J Nat Prod 64: 1286–1293

[8] Zheng ZP, Cheng KW, Chao J et al (2008) Tyrosinase inhibitors from paper mulberry (*Broussonetia papyrifera*). Food Chem 106: 529–535

[9] Ryu HW, Lee BW, Curtis-Long MJ et al (2010) Polyphenols from *Broussonetia papyrifera* displaying potent α-Glucosidase inhibition. J Agric Food Chem 58: 202–208

[10] Ryu HW, Curtis-Long MJ, Jung S et al (2012) Anticholinesterase potential of flavonols from paper mulberry (*Broussonetia papyrifera*) and their kinetic studies. Food Chem 132: 1244–1250

[11] Hubbe MA, Bowden C (2009) Handmade paper: a review of its history, craft, and science. BioRes 4(4): 1736–1792

[12] Qu LJ, Tian MW, Guo XQ et al (2014) Preparation and properties of natural cellulose fibres from *Broussonetia papyrifera* (L.) Vent Bast Fibres Text East Eur 22: 24–28

[13] Douglas AE (2006) Phloem-sap feeding by animals: problems and solutions. J Exp Bot 57(4): 747–754

[14] Kehr J (2006) Phloem sap proteins: their identities and potential roles in the interaction between plants and phloem-feeding insects. J Exp Bot 57(4): 767–774

[15] Turgeon R, Wolf S (2009) Phloem transport: cellular pathways and molecular trafficking. Ann. Rev Plant Biol 60: 207–221

[16] Scartazza A, Moscatello S, Matteucci G et al (2015) Combining stable isotope and carbohydrate analyses in phloem sap and fine roots to study seasonal changes of source-sink relationships in a Mediterranean beech forest. Tree Physiol 35: 829–839

[17] Nicolas S, Steinhauser D, Strelkov S et al (2005) GC-MS libraries for the rapid identification of metabolites in complex biological samples. FEBS Lett 579: 1332–1337

[18] Andersson-Gunnerås S, Mellerowicz EJ, Love J et al (2006) Biosynthesis of cellulose-enriched tension wood in *Populus*: Global analysis of transcripts and metabolites identified biochemical and developmental regulators in secondary wall biosynthesis. The Plant J 45(2): 144–165

[19] Lisec J, Schauer N, Kopka J et al (2006) Gas chromatography mass spectrometry-based metabolite profiling in plants. Nat Protoc 1: 387–396

[20] Yeh TF, Morris CR, Goldfarb B et al (2006) Utilization of polar metabolite profiling in the comparison of juvenile wood and compression wood in loblolly pine (*Pinus taeda*). Tree Physiol 26: 1497–1503

[21] Shi JT, Li J (2012) Metabolites and chemical group changes in the wood-forming tissue of *Pinus koraiensis* under inclined conditions. BioRe 7(3): 3463–3475

[22] IAWA Committee (2016) IAWA list of microscopic bark features. IAWA 37(4): 517–615

[23] Evert RF, Murmanis L (1965) Ultrastructure of the secondary phloem of Tilia. Am J Bot 52: 95–106

[24] Tucker CM, Evert RF (1969) Seasonal development of the secondary phloem in Acer negundo. Am J Bot 56: 275–284

[25] Antonova GF, Stasova VV (2006) Seasonal development of phloem in Scots pine stems. Russ J Dev Biol 37: 306–320

[26] Gričar J, Čufar K (2008) Seasonal dynamics of phloem formation in Silver fir and Norway spruce as affected by drought. Russ J Plant Physiol 55: 597–603

[27] Čufar K, Cherubini M, Gričar J et al (2011) Xylem and phloem formation in chestnut (*Castanea sativa* Mill.) during the 2008 growing season. Dendrochronologia 29: 127–134

[28] Jyske TM, Suuronen JP, Pranovich AV et al (2015) Seasonal variation in formation, structure, and chemical properties of phloem in *Picea abies* as studied by novel microtechniques. Planta 242: 613–629

[29] Dong M, Xu YM, Lin H et al (2016) Seasonal dynamics in cambial activity and the formation of xylem and phloem in the branches of *Cinnamomum camphora*. Dendrobiol 75: 13–21

[30] Esau K, Cheadle VI (1984) Anatomy of the secondary phloem of Winteraceae. IAWA Bull ns 5: 13–43

[31] Evert RF (2006) Esau’s Plant Anatomy: meristems, cells, and tissues of the plant body-their structure, function, and development. 3rd ed, John Wiley&Sons, Inc, New Jersey

[32] Villas-Boas SG, Noel S, Lane GA et al (2006) Extracellular metabolomics: a metabolic footprinting approach to assess fiber degradation in complex media. Analytical Biochem 349: 297–305

[33] Santos SAO, Freire CSR, Domingues MRM et al (2011) Characterization of phenolic components in polar extracts of *Eucalyptus globulus* L abill. Bark by high-performance liquid chromatography-mass spectrometry. J Agric Food Chem 59(17): 9386–9393

[34] Li X, Wu D, Lu T et al (2014) Highly efficient chemical process to convert mucic acid into adipic acid and DFT studies of the mechanism of the rhenium-catalyzed deoxydehydration. Angew Chem Int Edi 53(16): 4200

[35] Apelblat A (2014) Citric acid. Springer International Publishing, Berlin

[36] Fernie AR, Carrari F, Sweetlove LJ (2004) Respiratory metabolism: glycolysis, the TCA cycle and mitochondrial electron transport. Curr Opin Plant Biol 7: 254–261

[37] Babb VM, Haigler CH (2001) Sucrose phosphate synthase activity rises in correlation with high-rate cellulose synthesis in three heterotrophic systems. Plant Physiol 127: 1234–1242

[38] Pascual G, Avgustinova A, Mejetta S et al (2017) Targeting metastasis-initiating cells through the fatty acid receptor CD36. Nature 541: 41–45

[39] McLaurin J, Golomb R, Jurewica A et al (2000) Inositol stereoisomers stabilize an oligomeric aggregate of Alzehimer amyloid β-peptide and inhibit a β-induced toxicity. The J Biol Chem 275(24): 18495–18502

[40] Sanz ML, Villamiel M, Martínez-Castro I (2004) Inositols and carbohydrates in different fresh fruit juices. Food Chem 87: 325–328

[41] Bianchi S, Kroslakova I, Janzon R et al (2015) Characterization of condensed tannins and carbohydrates in hot water bark extracts of European softwood species. Phytochemistry 120: 53–61

[42] Chen J, Zhang HY, Chen YJ et al (2014) Optimization of D-pinitol extraction from vegetable soybean leaves and its potential application in control of cucumber powdery mildew. Crop Protection 60: 20–27

[43] Neri LM, Borgatti P, Tazzari PL et al (2003) The phosphoinositide 3-kinase/AKT1 pathway involvement in drug and all-trans-retinoic acid resistance of leukemia cells. Mol Cancer Res 1(3): 234–236

[44] Crabbe T (2007) Exploring the potential of PI3K inhibitors for inflammation and cancer. Biochem. Soc T 35(2): 253–256

[45] Wu P, Liu T, Hu Y (2009) PI3K inhibitors for cancer therapy: what has been achieved for far? Curr Med Chem 16(8): 916–930

[46] Kurtz JE, Ray-Coquard I (2012) PI3 kinase inhibitors in the clinic: an update, review. Anticancer Res 32(7): 2463–2470

[47] Onakpoya IJ, Spencer EA, Thompson MJ et al (2014) The effect of chlorogenic acid on blood pressure: a systematic review and meta-analysis of randomized clinical trials. J Hum Hypertens 29(2): 77–81

[48] Tajik N, Tajik M, Mack I et al (2017) The potential effects of chlorogenic acid, the main phenolic components in coffee, on health: a comprehensive review of the literature. Eur J Nutrition 56: 2215–2244

[49] Vila R, Mundina M, Tomi FL et al (2002) Composition and antifungal activity of the essential oil of *Solidago chilensis*. Planta Med 68(2): 164–167

[50] Lenfeld J, Kroutil M, Marsalek E et al (1981) Anti-inflammatory activity of quaternary benzophenanthridine alkaloids from *chelidonium majus*. J Med Plants Res 43: 161–165

